# GATA-1-dependent histone H3K27ac mediates erythroid cell-specific interaction between CTCF sites

**DOI:** 10.1101/2020.05.14.095547

**Authors:** Yea Woon Kim, Yujin Kang, Jin Kang, AeRi Kim

## Abstract

CTCF sites interact with each other in the chromatin environment, establishing domains. Our previous study showed that interaction between CTCF sites is cell type-specific around the β-globin locus and is dependent on erythroid specific activator GATA-1. To find out molecular mechanisms of the cell type-specific interaction, we directly inhibited GATA-1 binding to the β-globin enhancers by deleting its binding motifs and found that histone H3K27 acetylation (H3K27ac) was decreased at CTCF sites surrounding the β-globin locus, even though CTCF binding itself was maintained at the sites. Forced H3K27ac by TSA treatment or CBP/p300 KD affected the interactions between CTCF sites around the β-globin locus. Analysis of public ChIA-PET data revealed that H3K27ac is higher at CTCF sites forming short chromatin interactions than long interactions. The short interactions contain erythropoiesis-associated genes and GATA-1 binding sites in erythroid K562 cells. Depletion of GATA-1 reduced H3K27ac at CTCF sites near erythroid enhancers. These results indicate that GATA-1-dependent histone H3K27ac at neighboring CTCF sites mediates erythroid specific chromatin interaction between them.

## Introduction

Eukaryotic genomes are organized into three-dimensional structure by chromatin interactions. Close positioning between transcription regulatory regions, such as enhancers and promoters, has been well documented in many gene loci by chromosome conformation capture (3C) assay (Kim *et al*, 2009; Lee *et al*, 2017; Nativio *et al*, 2009; Palstra *et al*, 2003; Tolhuis *et al*, 2002). Genome-wide studies using high-throughput chromosome conformation capture (Hi-C) or chromosome conformation capture carbon-copy (5C) have revealed chromatin interaction domains of mega base scale, which are referred to as topologically associating domains (TADs) (Dixon *et al*, 2012; Nora *et al*, 2012). TADs insulate regions of large size for appropriate expression of genes (Dowen *et al*, 2014) and their boundaries are enriched with CCCTC-binding factor (CTCF), a well-known insulator protein (Dixon *et al.,* 2012; Rao *et al*, 2014). The association of CTCF protein is essential for forming chromatin domains including TADs as demonstrated by the depletion of CTCF protein or the deletion of its binding sites (Despang *et al*, 2019; Guo *et al*, 2015; Hou *et al*, 2010; Lee *et al*, 2019; Nora *et al*, 2017).

CTCF binding sites (CTCF sites) are scattered throughout the genome (Kim *et al*, 2007) and participate in most of chromatin interactions on Hi-C contact maps (Rao *et al.,* 2014). A small portion of CTCF sites is located in the boundary regions of TADs (Dixon *et al.,* 2012). CTCF-based intra-TADs or sub-TADs are observed inside of TADs (Matthews & Waxman, 2018). While TADs are large domains with ~1 Mb of median size and have relatively constant boundaries across different cell types (Dixon *et al.,* 2012), the enclosed sub-TADs are often cell type-specific (Brown *et al*, 2018; Hanssen *et al*, 2017; Matthews & Waxman, 2018). In the TAD containing the mouse α-globin genes and their regulatory elements, sub-TADs were formed in an erythroid specific fashion. However, the CTCF binding pattern through the TAD was similar in erythroid cells and non-erythroid cells (Brown *et al.,* 2018; Hanssen *et al.,* 2017). This result suggests that CTCF binding itself is not enough for forming chromatin interactions including tissue/cell type-specific sub-TADs.

The spatial organization of chromatin has been well characterized in the human β-globin locus. The β-globin locus contains locus control region (LCR) enhancer and the β-like globin genes that are specifically activated in erythroid cells. Five CTCF sites exist around the β-globin locus (Fig 1A). The C3 and C7 sites on both ends interact with each other, establishing a stable β-globin TAD, and other CTCF sites form erythroid specific sub-TADs (Huang *et al*, 2017; Kang *et al*, 2017). Our previous study showed that interactions between CTCF sites corresponding to sub-TADs are dependent on erythroid specific activator GATA-1 that binds to the LCR hypersensitive sites (HSs) (Kang *et al.,* 2017). GATA-1 mediates chromatin interaction between enhancers and promoters in the β-globin locus (Kim *et al*, 2011) and recruits co-activators such as histone acetyltransferase CBP and p300 to enhancers (Kiekhaefer *et al*, 2002; Letting *et al*, 2003). In addition, GATA-1 acts as a master regulator of erythropoiesis, activating the genes encoding biosynthetic enzymes and diverse red cell constituents (Tanimura *et al*, 2016).

**Figure 1.**
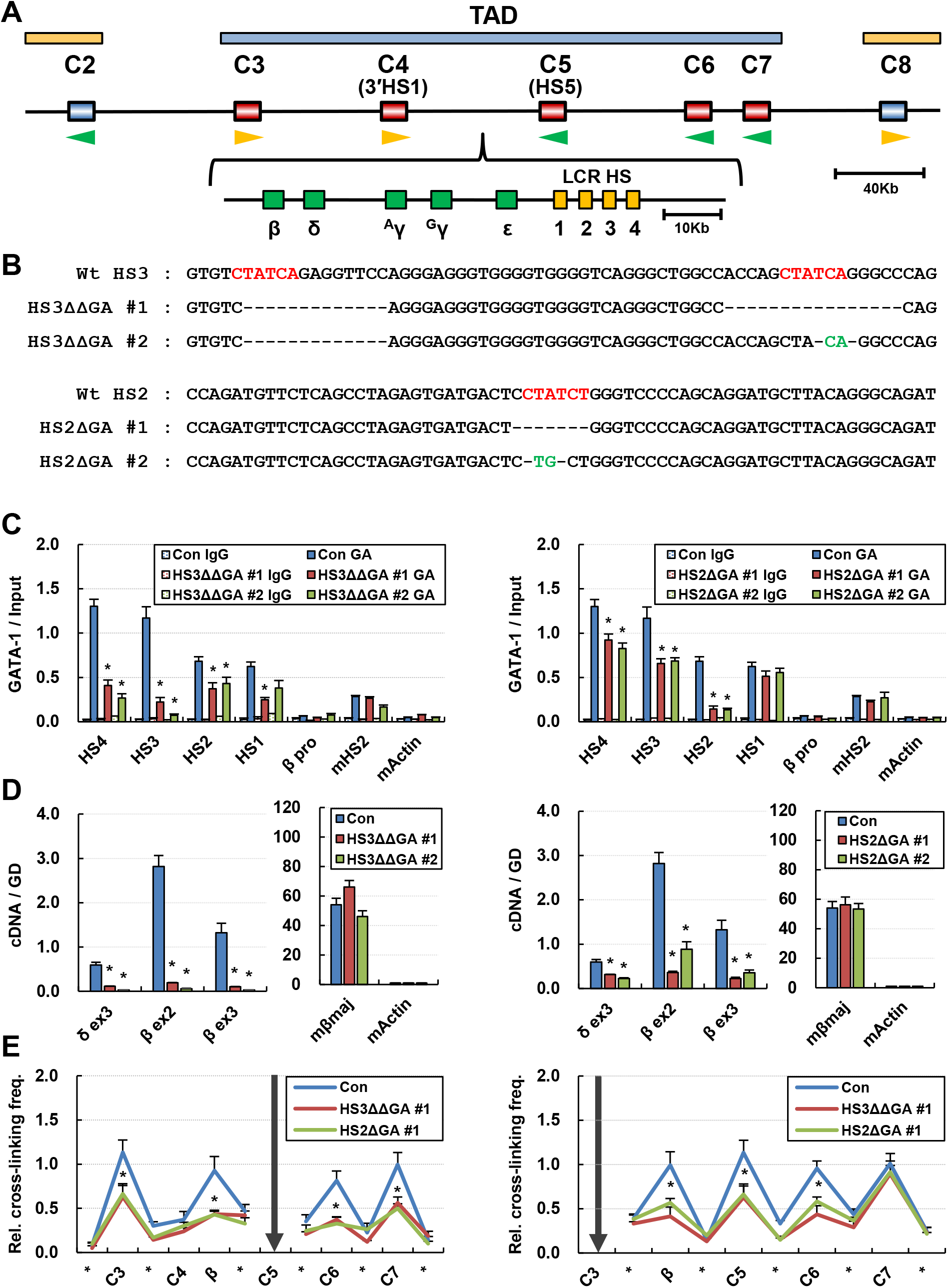
Chromatin interaction between CTCF sites around the β-globin locus having deletion of GATA-1 binding motifs. (A) CTCF sites are indicated around the human β-globin locus by squares and are named as C2-C8. Red squares are CTCF sites inside of the β-globin TAD and blue squares are outside of the TAD. The β-globin locus is located between C4 and C5. (B) GATA-1 binding motifs were deleted from the β-globin LCR HS3 and HS2 by CRISPR/Cas9 system in MEL/ch11 cells. DNA sequences for the LCR HS3 and HS2 are presented in wild type cells (Wt) and GATA-1 motif deleted clones (HS3ΔΔGA#1, 2 and HS2ΔGA#1, 2). GATA-1 binding motifs are marked by red color. Black dashes are nucleotides deleted and green bases are nucleotides replaced. (C) ChIP was performed with GATA-1 antibodies in control cells (Con) and GATA-1 motifs deleted clones (left; HS3ΔΔGA and right; HS2ΔGA). Amounts of immunoprecipitated DNA were quantitatively compared with amounts of input DNA for the indicated PCR amplicons. The mouse Actin gene served as an internal negative control and normal IgG (IgG) served as an experimental negative control. (D) Transcription levels of the human δ- and β-globin genes and mouse β major globin gene were determined by quantitative RT-PCR in control cells (Con) and GATA-1 motifs deleted clones (left; HS3ΔΔGA and right; HS2ΔGA). Amounts of cDNA for target genes were compared with a genomic DNA standard and then normalized to the mouse Actin cDNA. (E) 3C assay was performed in control cells (Con) and GATA-1 motifs deleted clones (HS3ΔΔGA#1 and HS2ΔGA#1). Fragments containing the C5 or the C3 were used as anchors to measure relative cross-linking frequencies with fragments containing CTCF sites or fragments not containing them. The fragments not containing a CTCF site are represented by asterisks on the x-axis between CTCF sites. Relative cross-linking frequencies were determined by quantitatively comparing ligated DNA in cross-linked chromatin with control DNA and then by normalizing to the cross-linking frequency in the mouse Ercc3 gene. The results are the means of five to six independent experiments ± SEM. *P < 0.05.

Here, to explore molecular mechanisms of GATA-1-dependent interaction between CTCF sites, we directly inhibited GATA-1 binding to the β-globin LCR HS3 and HS2 by deleting its binding motifs, and examined interactions between neighboring CTCF sites and chromatin structure at the CTCF sites. Based on the reduction of H3K27ac at CTCF sites by the loss of GATA-1 binding, the effects of this histone modification were investigated using histone deacetylase inhibitor TSA-treated cells and histone acetyltransferase CBP/p300 KD cells. The relationship between H3K27ac and chromatin interaction was analyzed using public ChIA-PET data. Finally, we analyzed the role of GATA-1 in H3K27ac at CTCF sites near erythroid specific enhancers. Our studies showed that tissue specific activator GATA-1 plays a role in H3K27ac at neighboring CTCF sites in erythroid cells and this histone modification is required for cell type-specific chromatin interactions between CTCF sites.

## Results

### Loss of GATA-1 binding at the LCR HS3 or HS2 decreases interactions between CTCF sites around the β-globin locus

To ask whether GATA-1 directly affects interaction between CTCF sites around the human β-globin locus, we deleted GATA-1 binding motifs at the human LCR HS3 or HS2 enhancer in erythroid MEL/ch11 cells containing a human chromosome 11. Two GATA-1 motifs were deleted at the LCR HS3 by two kinds of guide RNAs, respectively, with CRISPR/spCas9 system (Fig 1B), resulting in severe inhibition of GATA-1 binding (Fig 1C). GATA-1 binding to the HS2 was impaired by the deletion of one motif for it. The inhibition of GATA-1 binding to the HS3 or HS2 affected its binding to the other LCR HSs. It might be due to physical and functional interaction between the HSs in the erythroid specific chromatin environment. This GATA-1 binding inhibition resulted in transcriptional failure of the β-globin gene (Fig 1D). Consistent results were obtained for two independent clones, #1 and # 2, having GATA-1 motif deletion at the HS3 or HS2 (HS3ΔΔGA or HS2ΔGA).

Next, we analyzed chromatin interactions between five CTCF sites (C3-C7) around the β-globin locus using the 3C assay. The loss of GATA-1 binding decreased cross-linking frequency between the C5 anchor fragment and those containing C3, C6, C7 or the β-globin gene (Fig 1E). When fragments containing C3 were used as an anchor, they cross-linked with C7 at similar frequency in HS3ΔΔGA#1, HS2ΔGA#1 and control cells, but their crosslinking with C5, C6 and β-globin gene was reduced in the GATA-1 motif deletion clones. These results indicate that GATA-1 contributes to interaction between CTCF sites around the β-globin locus. When it is compared to results from knocking down GATA-1, results from deleting its motifs support more direct roles of GATA-1. However, chromatin interaction corresponding to the β-globin TAD appears to be unaffected by the loss of GATA-1 binding.

### CTCF and Rad2l are recruited to CTCF sites around the β-globin locus without LCR GATA-1 binding, but H3K27ac is reduced at the CTCF sites

To explore the molecular mechanism of GATA-1-dependent chromatin interaction between CTCF sites, we examined chromatin structure at CTCF sites around the β-globin locus. The 5 CTCF sites were occupied by CTCF and Rad21 as shown by ChIP assay, even the occupancy levels at C4, C5 and C6 were lower than the levels at C3 and C7 but higher than the level at negative region mActin. The loss of LCR GATA-1 binding did not affect the occupancy of CTCF and Rad21 in HS3ΔΔGA#1 and HS2ΔGA#1 clones (Fig 2A, B). Histone H3 depletion was maintained at the CTCF sites without HS3 or HS2 GATA-1 binding (Fig 2C). However, we found that histone H3K27 acetylation (H3K27ac), an active histone modification, is reduced at the CTCF sites in the GATA-1 motif deletion clones (Fig 2D). Extended analysis using ChIP-seq showed that the CTCF sites around the β-globin locus are hyperacetylated at H3K27 and this modification is decreased by the loss of GATA-1 binding to the LCR HS3 (Fig 2E). H3K27ac reduction was not observed in the regions flanking the β-globin TAD (Fig S1). Taken together, GATA-1 occupancy in the LCR appears to have a role in establishing histone H3K27ac at CTCF sites around the β-globin locus. CTCF binding itself does not appear to correlate with H3K27ac at the sites.

**Figure 2.**
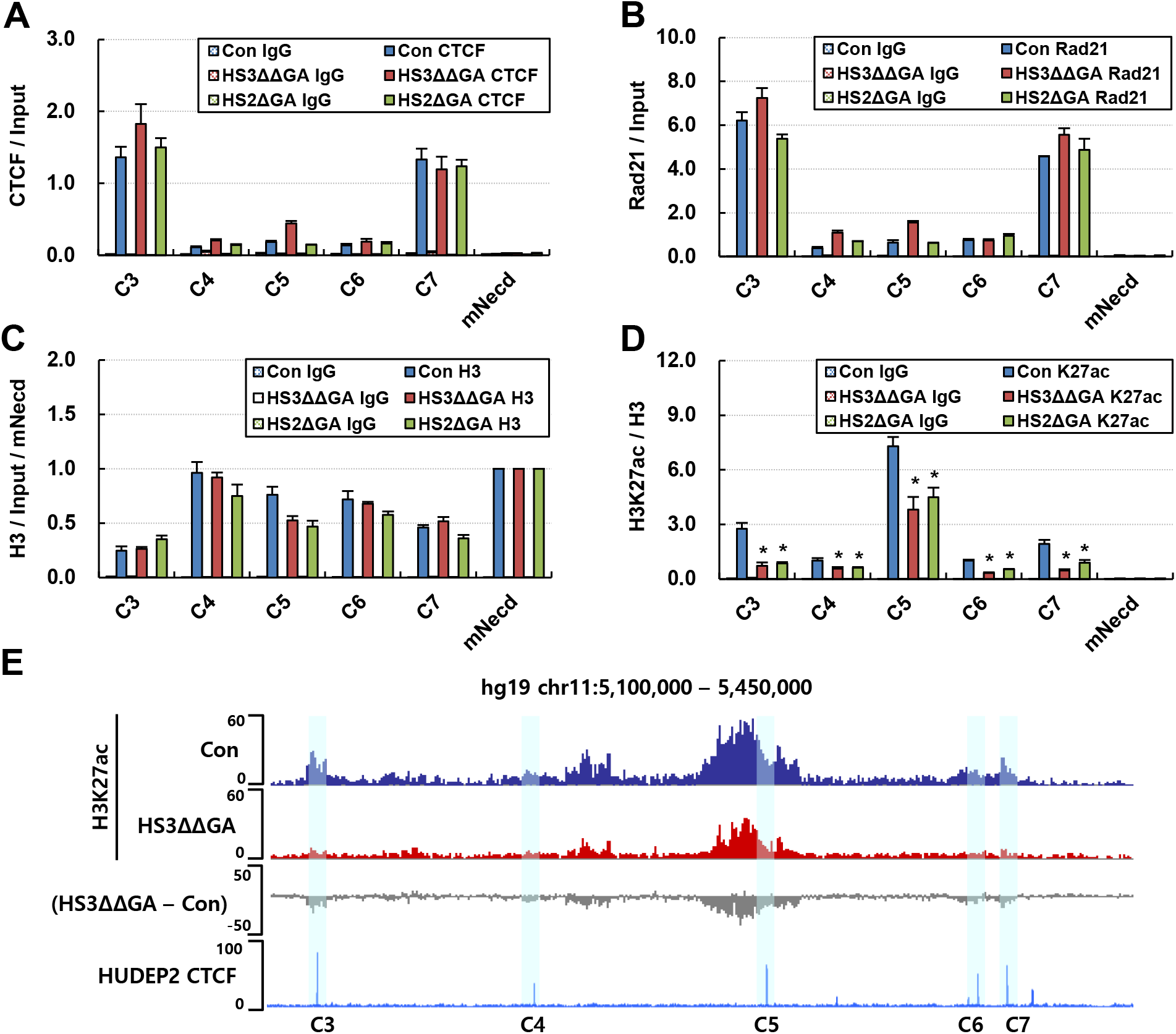
Chromatin structures at CTCF sites around the β-globin locus having deletion of GATA-1 binding motifs. ChIP was performed with antibodies specific to CTCF (A), Rad21 (B), histone H3 (C) and H3K27ac (D) in control cells (Con) and HS3ΔΔGA and HS2ΔGA clones. Amounts of immunoprecipitated DNA were determined as described in Fig 1C. The mouse Necdin gene served as an internal control. The results are the means of four independent experiments ± SEM. *P < 0.05. (E) Genome browser for ChIP-seq data presents the distribution of H3K27ac around the β-globin locus. Difference of H3K27ac between control and HS3ΔΔGA (HS3ΔΔGA - Con) is presented by gray track. CTCF sites are highlighted with blue shadows using public CTCF ChIP-seq data from HUDEP2 cells where human adult β-globin gene is predominantly transcribed.

### Increase of H3K27ac induces chromatin interactions between CTCF sites around the β-globin locus

Results from the β-globin locus where LCR GATA-1 binding is inhibited suggested that histone H3K27ac can mediate interaction between CTCF sites. To examine this possibility, we induced H3K27ac by treating MEL/ch11 cells with histone deacetylase inhibitor Trichostatin A (TSA). TSA treatment for 6 h and 24 h elevated H3K27ac at a total protein level as shown by western blot (Fig 3A). H3K27ac at a chromatin level was increased at the 5 CTCF sites around the β-globin locus as measured by ChIP assay (Fig 3B). The forced H3K27ac increased interaction of C5 with C3 and C7, and interaction of C3 with C6 (Fig 3C). However, interaction between the C3 and C7 corresponding to TAD borders was unaffected, and CTCF and Rad21 occupancies were maintained at the 5 CTCF sites (Fig 3D, E). Thus these results suggest that H3K27ac plays a role in enhancing interaction between CTCF sites to form sub-TADs. The binding of CTCF and Rad21, itself, appears to be not enough for interaction between CTCF sites.

**Figure 3.**
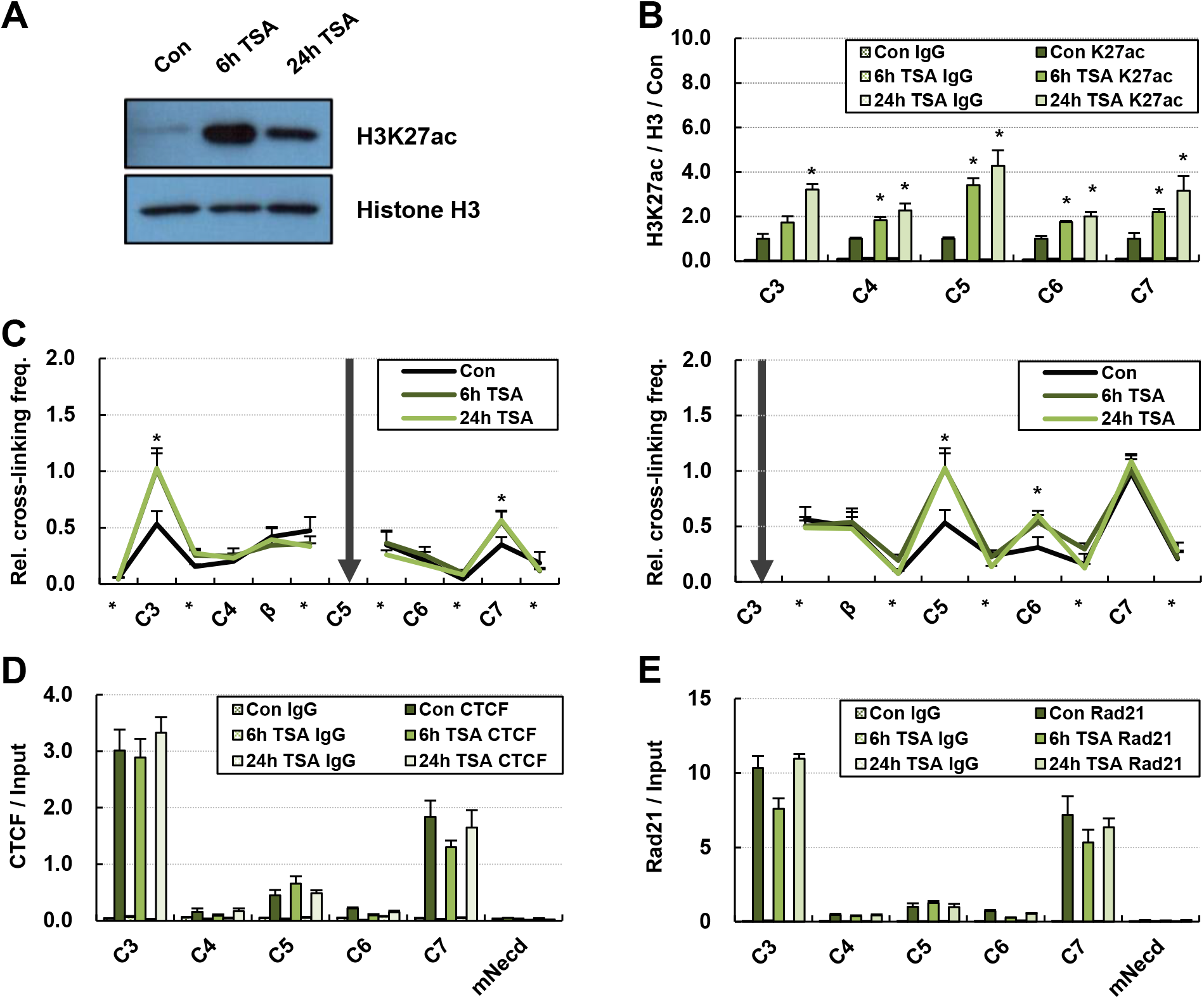
Chromatin interaction between CTCF sites around the β-globin locus in TSA treated MEL/ch11 cells. (A) MEL/ch11 cells were treated with 25 ng/ml of TSA for 6 h or 24 h. Histone H3 acetylated at K27 was detected by Western blotting in nuclear extract from control cells (Con) and cells treated with TSA. Histone H3 was used as an experimental control. (B) H3K27ac was determined in CTCF sites around the β-globin locus by ChIP. DNA immunoprecipitated by H3K27ac antibodies were quantitatively compared with DNA immunoprecipitated by H3, and then normalized to value in control cells. Normal IgG (IgG) was used as an experimental negative control. (C) Relative cross-linking frequencies was determined between CTCF sites around the β-globin locus in 3C assay as described in Fig 1E. Fragments containing C5 and C3 were used as anchors. Occupancies of CTCF (D) and Rad21 (E) were determined at the CTCF sites by ChIP. Results are presented as the means ± SEM of four to six independent experiments in ChIP and 3C assay. *P < 0.05.

### Decrease of H3K27ac does not affect genome-wide CTCF binding, but disrupts chromatin interactions between CTCF sites around the β-globin locus

To further study the effect of histone H3K27ac on genome-wide CTCF binding and chromatin interaction between CTCF sites, we inhibited the expression of histone acetyltransferase CBP and p300 in human erythroid K562 cells (C/pi) and analyzed the change of H3K27ac using ChIP-seq approach. Our H3K27ac ChIP-seq data from control cells were consistent with public ChIP-seq data from K562 cells (Fig S2). H3K27ac was decreased at 84.7 % of its peaks (n=83,364) by the CBP/p300 knockdown (Fig 4A). When total CTCF sites were sorted by H3K27ac level in wild type K562 cells and control K562 cells, a portion of them was highly acetylated at H3K27 (Fig 4B). CTCF binding did not correlate with H3K27ac in either wild type or control K562 cells. To precisely analyze the role of H3K27ac in CTCF binding, we divided hyperacetylated CTCF sites (top 50%, n=28,586) into two groups; one group where H3K27ac was decreased (58%, n=16,693) and the other group where it was unchanged or increased by C/pi (Fig 4C). Both groups showed unchanged CTCF binding, indicating that histone H3K27ac is not necessary for CTCF binding. However, this was not the case for chromatin interactions between CTCF sites. Reduction of H3K27ac disrupted interaction between CTCF sites around the β-globin locus, even though CTCF binding was maintained (Fig 4D, E). Taken together, while H3K27ac is not required for CTCF binding, it may be important for chromatin interaction between CTCF sites.

**Figure 4.**
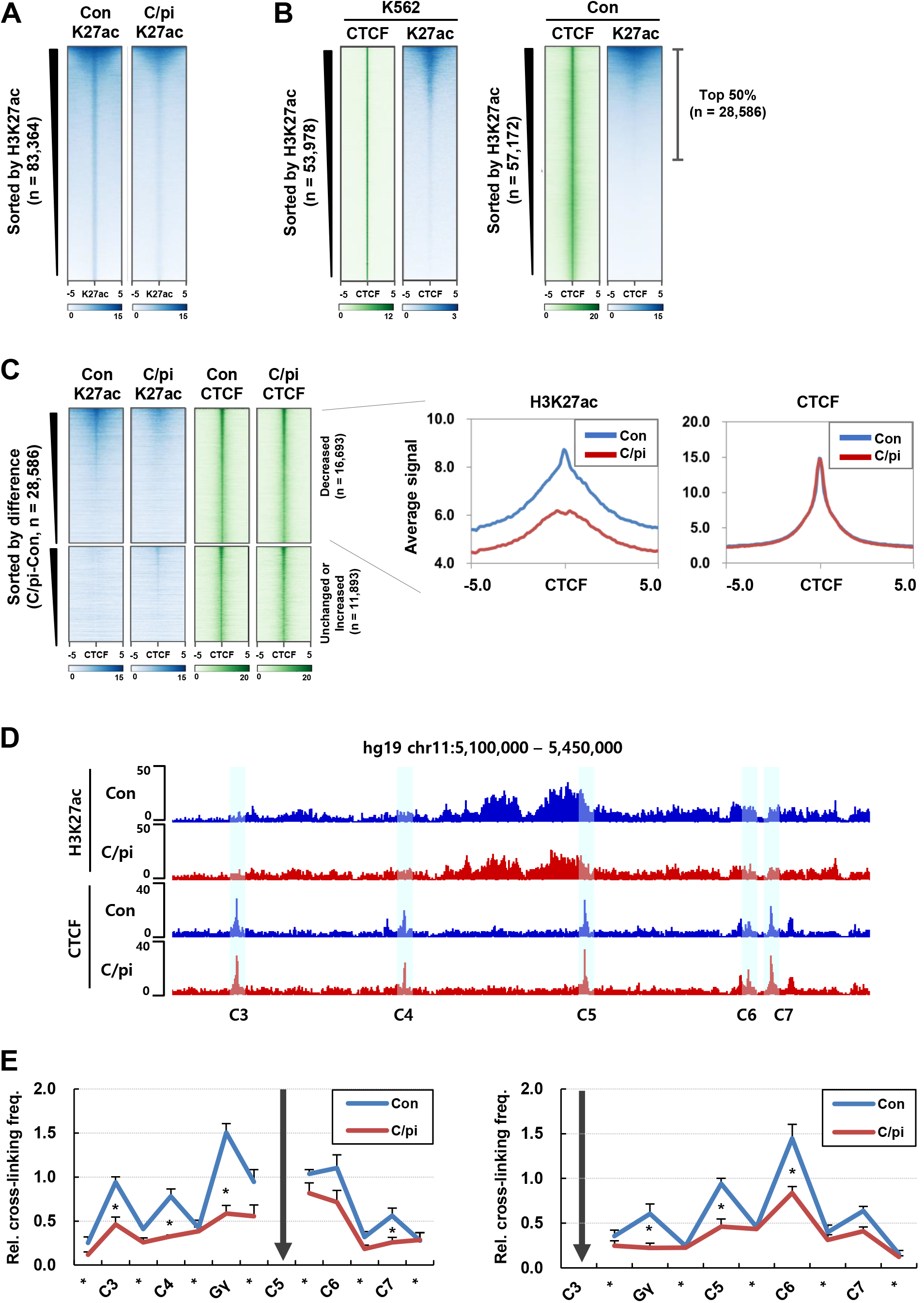
CTCF binding and chromatin interaction in CBP/p300 knockdown cells. ChIP-seq for CTCF and H3K27ac was performed in control cells (Con) and CBP/p300 knockdown K562 cells (C/pi). (A) Heatmaps represent H3K27ac ChIP-seq signals at its peaks (n=83,364) in control cells (Con) and CBP/p300 knockdown cells (C/pi). (B) CTCF peaks were sorted by H3K27ac in control K562 cells and wild type K562 cells using public ChIP-seq data. (C) Top 50 % of CTCF sites (n=28, 568) was divided into two groups according to changes of H3K27ac by CBP/p300 KD; decreased one (0 > C/pi CPM - Con CPM) and unchanged or increased one (0 ≤ C/pi CPM - Con CPM). CTCF signals were aligned in heatmaps. Average profiles of H3K27ac and CTCF signal were drawn for a group where H3K27ac is decreased in CBP/p300 knockdown cells. (D) The β-globin locus was presented with view of genome browser for ChIP-seq data in control cells and CBP/p300 knockdown cells. CTCF sites were highlighted with blue shadows. (E) Relative cross-linking frequencies were determined in CTCF sites around the β-globin locus by 3C analysis as described in Fig 1. Results are presented as the means ± SEM of five independent experiments. *P < 0.05.

### CTCF sites forming short interactions are highly acetylated at H3K27 and erythroid specific genes are present in the short interactions in K562 cells

CTCF sites participating in chromatin interaction can be identified using paired-end tag sequencing (ChIA-PET) data that shows chromatin interactions at regions bound by target protein. To study relationship between H3K27ac at CTCF sites and chromatin interaction, we analyzed public CTCF ChIA-PET data and ChIP-seq data for CTCF and H3K27ac that were generated using K562 cells. 25,721 CTCF-associated chromatin interactions were identified from the ChIA-PET data that include 834 inter-chromosomal interactions. The distance of CTCF-associated intra-chromosomal interactions (n=24,887) was computationally normalized to 100 Kb, except for 2 Kb at boundaries of the interactions. The 20 Kb regions flanking both sides of the interactions were included in the analysis. As expected, analysis using public ChIP-seq data showed that CTCF is enriched in the boundaries of CTCF-associated interactions (Fig 5A). H3K27ac was also enriched in the boundaries. When the CTCF-associated interactions were sorted by their size, H3K27ac levels were decreased according to the increase of interaction size (Fig 5B). The boundaries (anchors) of relatively short interactions (less than 100 Kb, n=9,287) were highly acetylated at H3K27ac, but the acetylation level was decreased remarkably in the boundaries of longer interactions (over than 100 Kb) (Fig 5C). CTCF binding did not show any correlation with the size of interactions. GO biological process analysis showed that the short CTCF-associated interactions contain genes (n=5,014) encoding mainly proteins for zinc ion homeostasis and cellular response to copper ion that are essential for erythropoiesis (Fig 5D). Erythroid specific activator GATA-1 was identified as a transcription factor that mostly relates with these genes. Indeed, GATA-1 binding peaks were present in 40 % of the short CTCF-associated interactions (Fig 5E). These results suggest that H3K27ac at the CTCF sites relates closely to short CTCF-associated chromatin interactions. A large portion of short interactions might be erythroid specific in K562 cells.

**Figure 5.**
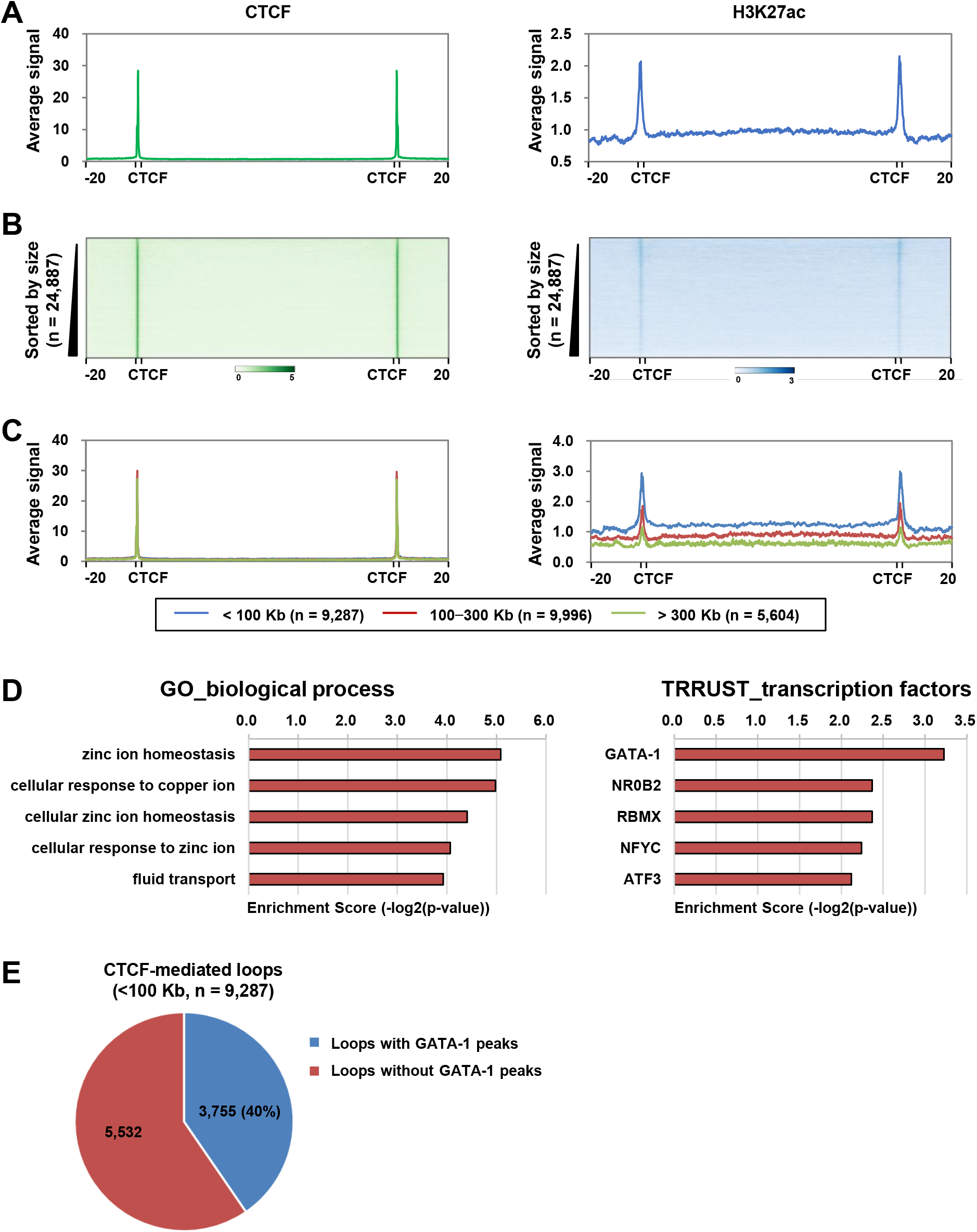
Analysis of relationship between H3K27ac and chromatin interaction using CTCF ChIA-PET data. (A) Average profiles of CTCF and H3K27ac signal were drawn in CTCF-mediated chromatin interactions using public CTCF ChIA-PET data in K562 cells. (B) Chromatin interactions were sorted by size, and heatmaps of CTCF and H3K27ac ChIP-seq signals were drawn in the interactions (C) Average profiles of CTCF and H3K27ac signal were drawn depending on size of CTCF-mediated chromatin interactions. (D) GO biological process analysis and TRRUST_transcription factor analysis were performed in CTCF-mediated chromatin interactions less than 100 Kb. (E) CTCF-mediated chromatin interactions less than 100 Kb were divided depending on presence of GATA-1 binding peak in a pie chart.

### GATA-1 depletion decreases H3K27ac at CTCF sites near erythroid specific enhancers

To further study the role of GATA-1 in H3K27ac at neighboring CTCF sites, we knocked down GATA-1 expression in K562 cells (Kim *et al.,* 2011) and carried out H3K27ac ChIP-seq. CTCF sites surrounding the β-globin locus showed that H3K27ac levels are decreased by the GATA-1 depletion (Fig 6A). To analyze the role of GATA-1 at a genome-wide level we defined erythroid specific enhancers (n=6,860) by overlapping peak regions of GATA-1, erythroid activator TAL1 and p300 in public ChIP-seq data (Fig 6B). While H3K27ac at total CTCF sites (n=57,172) was largely maintained in GATA-1 KD cells compared to control cells (Fig 6C), the CTCF sites (n=4,440) within 20 Kb from the erythroid enhancers showed the reduction of H3K27ac upon the GATA-1 depletion (Fig 6D). These results imply that the role of GATA-1 in H3K27ac at neighboring CTCF sites is not limited to the β-globin locus but can be applied genome-widely in erythroid cells.

**Figure 6.**
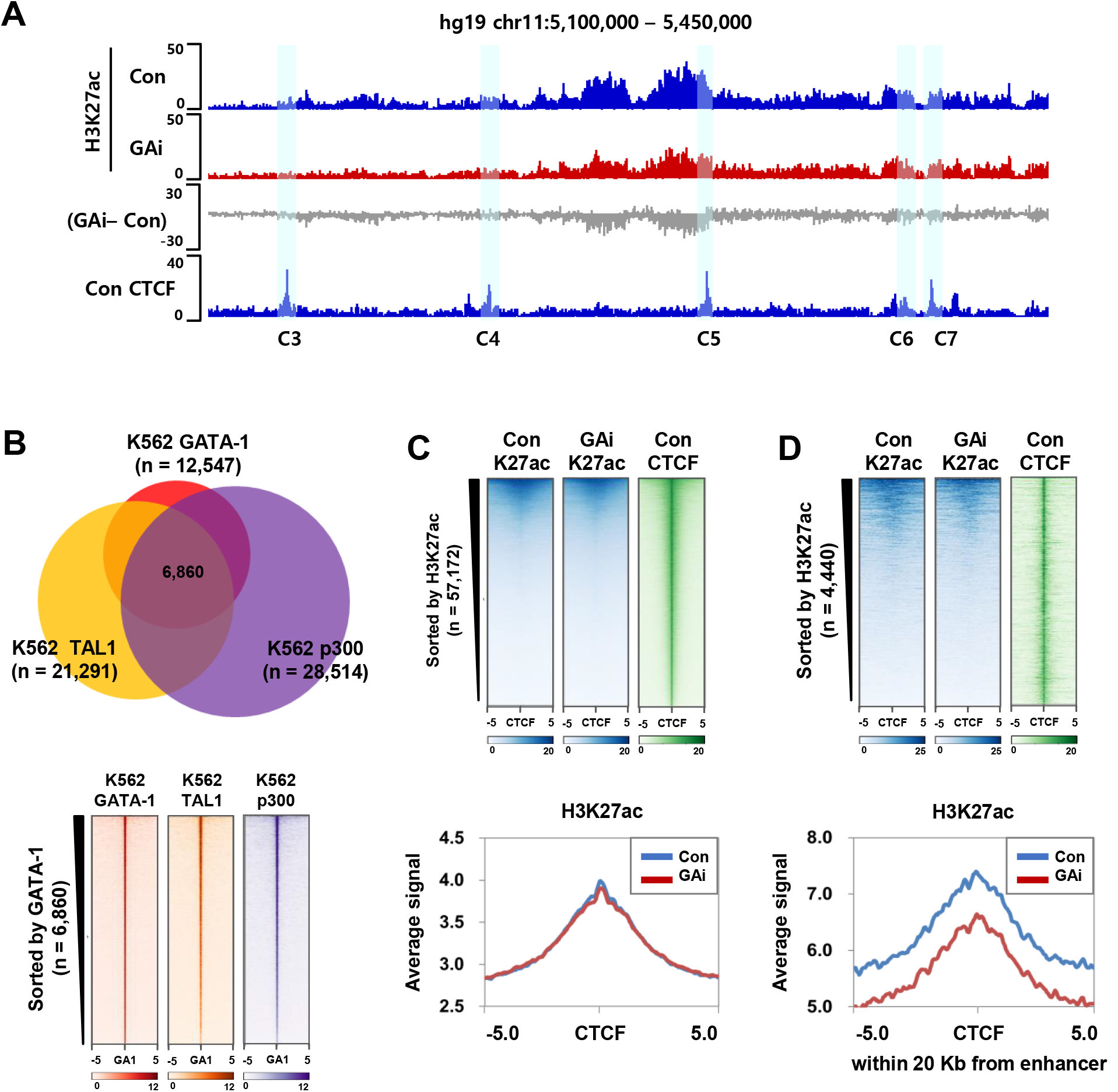
H3K27ac at CTCF sites around erythroid specific enhancers in GATA-1 knockdown K562 cells. (A) Genome browser for ChIP-seq data presents the distribution of H3K27ac around the β-globin locus. Difference of H3K27ac between control and GATA-1 knockdown (GAi - Con) is presented by gray track. CTCF sites were highlighted with blue shadows. (B) GATA-1 binding sites were overlapped with TAL1 and p300 sites in a venn diagram and heatmap to define putative erythroid specific enhancers. (C) Total CTCF peaks (n=57,172) were sorted by H3K27ac in control cells (Con), and H3K27ac signal at the peaks was presented by heatmaps and average profiles with it in GATA-1 knockdown cells (GAi). (D) CTCF sites were identified within 20 Kb of erythroid specific enhancers in control cells (n=4,440), and H3K27ac signal at the CTCF sites was presented by heatmaps and average profiles with it in GATA-1 knockdown cells.

## Discussion

This study shows that erythroid specific activator GATA-1 binding to the β-globin LCR HSs is required for histone H3K27ac at neighboring CTCF sites, which contributes to cell typespecific chromatin interaction between the CTCF sites (Fig 7A). The positive role of histone H3K27ac in the chromatin interaction was demonstrated in TSA treated cells and CBP/p300 KD cells where this modification was forced and reduced, respectively. CTCF binding itself appears to be not enough for the chromatin interactions. Analysis according to the length of CTCF-associated interactions indicates that H3K27ac at CTCF sites associates with short and cell type-specific interactions such as sub-TADs. CTCF sites near erythroid specific enhancers were acetylated at H3K27 in a GATA-1-dependent manner. Therefore, we propose that tissue specific activator GATA-1 induces H3K27ac at neighboring CTCF sites in erythroid cells. Histone H3K27ac at CTCF sites appears to promote chromatin interactions that are tissue/cell type-specific.

**Figure 7.**
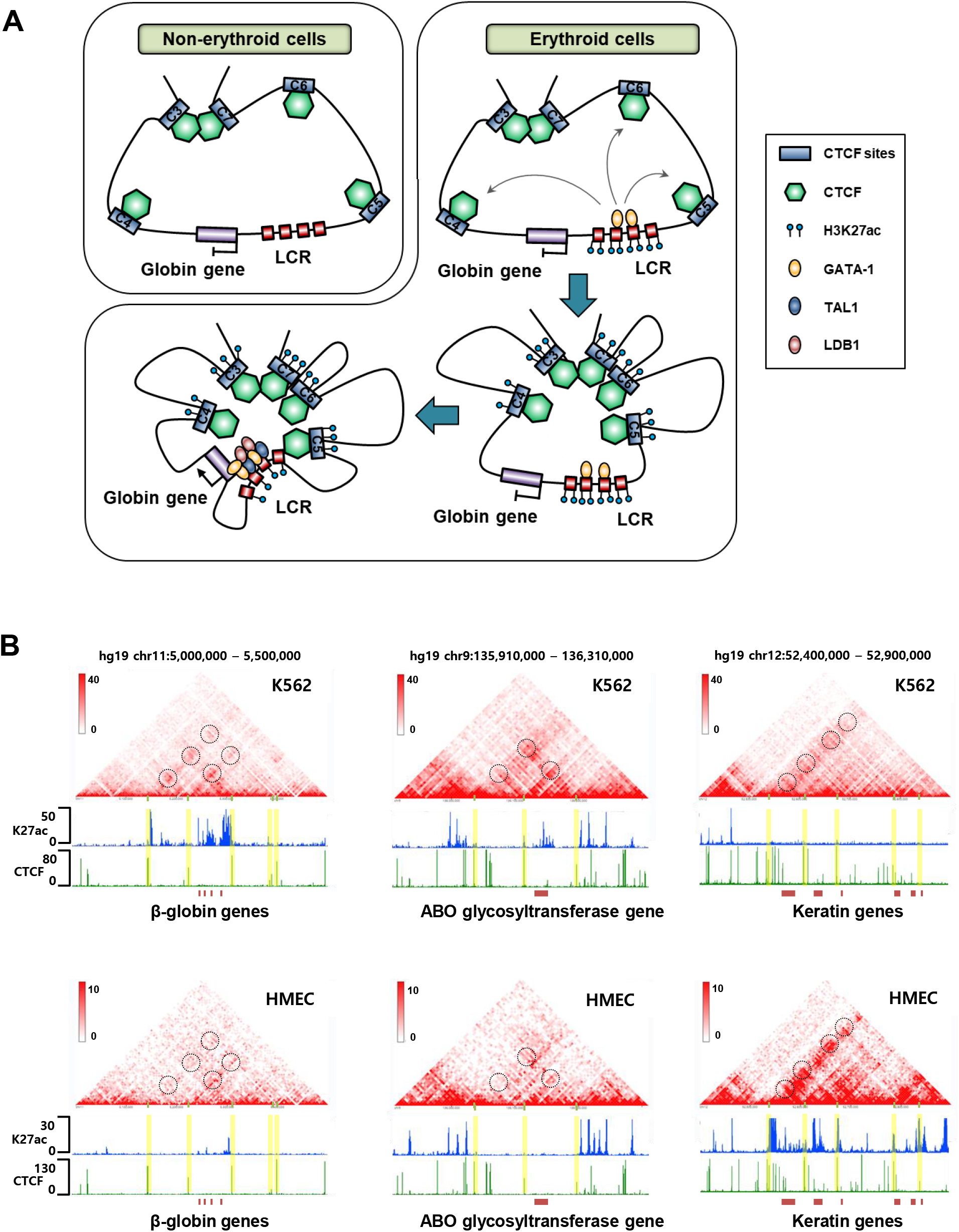
Chromatin structures around the β-globin locus. (A) The results from Fig 1–4 and our previous studies (Kang *et al.,* 2017; Yun *et al.,* 2014) were combined and represented for the human β-globin locus. CTCF binds to its motif in non-erythroid cells but does not induce interactions between the CTCF sites. In erythroid cells, GATA-1 mediates H3K27ac at neighboring CTCF sites, resulting in interactions between them. (B) Hi-C interactions, H3K27ac and CTCF ChIP-seq signals were aligned around the β-globin locus, the ABO glycosyltransferase gene and the Keratin genes in erythroid K562 cells (upper) and non-erythroid HMEC cells (lower). Yellow shadows and dashed circles indicate CTCF sites and chromatin interactions between CTCF sites, respectively.

GATA-1 has been known to establish histone acetylation on erythroid gene loci by recruiting CBP/p300 acetyltransferases and to be responsible for enhancer-promoter looping by interacting with LDB1 complex (Letting *et al.*, 2003; Song *et al*, 2007). This study shows that GATA-1 plays a role in histone H3K27ac at neighboring CTCF sites, contributing to chromatin interaction between CTCF sites surrounding the β-globin locus. The GATA-1-dependent H3K27ac is found at numerous CTCF sites near erythroid specific enhancers. However, GATA-1 binding sites rarely overlap with CTCF binding sites in K562 cells (Fig S3). Instead, cohesin complex consisting of Smc1, Smc3 and Rad21 subunits occupies both enhancers and CTCF sites (Kagey *et al*, 2010) and is found at the β-globin LCR HSs and CTCF sites in erythroid cells (Chien *et al*, 2011). The binding of Rad21 to the β-globin LCR HSs is dependent on GATA-1 in K562 cells (Kang *et al.,* 2017). Thus, GATA-1 could contribute to H3K27ac at CTCF sites by recruiting histone acetyltransferases and transferring them to CTCF sites using cohesin. In another possibility, enhancer-promoter interaction mediated by GATA-1 can affect CTCF sites around the β-globin locus. However interactions between the CTCF sites are maintained in another erythroid activator TAL1 KD cells where the enhancer-promoter interaction is disrupted (Kang *et al.,* 2017; Yun *et al*, 2014). Thus, GATA-1 might have a direct role in H3K27ac at neighboring CTCF sites, contributing to chromatin interaction between the CTCF sites. GATA-1-dependent interaction between CTCF sites has been found around the TAL1 locus (Zhou *et al*, 2013).

Histone modifications at CTCF binding sites have not been extensively studied because the depletion of CTCF protein or the deletion of its binding motifs resulted in dramatic effects on gene transcription and chromatin interaction. However, genome-wide analysis shows that a part of CTCF sites is highly acetylated at histone H3K27 in K562 cells. This pattern is observed in human epidermal keratinocytes (Rubin *et al*, 2017). Compared to the inside regions of CTCF-associated interactions in ChIA-PET analysis, the anchors are highly acetylated at histone H3K9 and H3K27 (Tang *et al*, 2015). A study using Hi-C technique shows a positive correlation between H3K27ac level and interaction density among regulatory regions (promoter, enhancer and insulator) (Ren *et al*, 2017). These reports support that H3K27ac acts favorably for chromatin interactions between CTCF sites, which is compatible with our results showing H3K27ac dependent-interactions between CTCF sites around the β-globin locus.

Even though we could not elucidate what is the direct role of H3K27ac at CTCF sites in chromatin interactions, one possible explanation is that this modification can be used to recruit coactivators such as BET family proteins (Filippakopoulos & Knapp, 2014). Recent studies show that BRD2, one of the BET proteins, co-localizes with CTCF and Rad21 and physically associates with them (Cheung *et al*, 2017; Hsu *et al*, 2017). The depletion of BRD2 disrupts the CTCF/BRD2 co-occupied boundaries of chromatin domains (Hsu *et al.,* 2017). BRD2 can form homodimers or heterodimers with another BET proteins (Garcia-Gutierrez *et al*, 2012; Nakamura *et al*, 2007). Thus H3K27ac at CTCF sites can have a role in recruiting coactivators to strengthen and stabilize interaction between CTCF sites.

H3K27ac at CTCF sites is likely to contribute to formation of cell-type specific sub-TADs rather than constant TADs. In the 3C analysis around the β-globin locus, interactions between CTCF sites corresponding to cell-type specific sub-TADs were affected by the changes of H3K27ac level. H3K27ac was greater at the anchors of CTCF-associated chromatin interactions less than 100 Kb in comparing with them of the longer chromatin interactions. Public ChIP-seq data and Hi-C data for K562 cells and HMEC mammary epithelial cells show that H3K27ac at the CTCF sites correlates with cell-type specific interactions between CTCF sites around the gene loci including the β-globin locus, ABO glycosyltransferase gene and Keratin gene (Fig 7B). However, CTCF binding patterns around the loci are similar in both cells, implying that CTCF binding itself is not enough for chromatin interactions. When chromatin interaction is disrupted around the β-globin locus by the loss of GATA-1 binding or CBP/p300 proteins, CTCF binding is unaffected around the locus. Unaffected CTCF binding is also observed at the boundaries of chromatin domains impaired by the depletion of BRD2 (Hsu *et al.,* 2017). Therefore, H3K27ac at CTCF sites is thought to be important for cell type specific sub-TAD formation. In erythroid cells, tissue specific activator GATA-1 appears to play a role in inducing H3K27ac at neighboring CTCF sites.

## Materials and Methods

### Cloning guide sequences of RNA into CRISPR vectors

GATA-1 binding motifs were deleted at the β-globin LCR HS3 and HS2 in MEL/ch11 cells by the CRISPR/spCas9 system as described previously (Kim & Kim, 2017). Sequences of oligonucleotides for single guide RNA (sgRNA) are presented in Supplementary Table S1. Two oligo F and R for sgRNA targeting GATA-1 motifs were phosphorylated, annealed and ligated into lentiCRISPRv2 vector (Addgene #52961) (Sanjana *et al*, 2014) for hHS3_GA oligo1 and hHS2_GA oligo or pLH-spsgRNA2 vector (Addgene #64114) (Ma *et al*, 2015) for hHS3_GA oligo2 as suggested by the manufacturer. Ligated plasmid DNA was transformed into competent Stbl3 bacteria and purified by Plasmid Midi kits (Qiagen).

### Cell culture and lentiviral transduction

MEL/ch11 and 293FT cells were cultured in DMEM medium containing 10% FBS. sgRNA-cloned CRISPR vector was co-transfected into 293FT cells with Virapower packaging mix using Lipofectamine 2000 (Invitrogen). At 72 h after transfection, supernatant was mixed with MEL/ch11 cells in the presence of 6 μg/ml polybrene. Cells were selected by 2 μg/ml puromycin for lentiCRISPRv2 or by 500 μg/ml hygromycin for pLH-spsgRNA2 at 72 h after transduction and clonal cells were isolated by dilution in a 96-well plate. Genomic DNA flanking the LCR HS3 or HS2 was amplified by PCR, and GATA-1 motif deletions were detected by sequencing the PCR products. For transcriptional activation of the β-globin gene, MEL/ch11 cells were induced at a concentration of 1.5 × 10^5^ cells/ml with 5 mM HMBA for 72 h.

### Chromatin immunoprecipitation (ChIP)

Cells (2 × 10^7^) were cross-linked with 1% formaldehyde at 25°C for 10 min. Nuclei were digested with 100 units of MNase at 37°C for 15 min. Mono or dinucleosomes-sized chromatin was incubated with antibodies for 3 h at 4°C and recovered with protein A agarose beads. DNA was purified by phenol extraction and then analyzed by quantitative PCR. The sequences of primers for ChIP assay are presented in Supplementary Table S2 and S3. Antibodies used in ChIP experiment are GATA-1 (sc-1233) from Santa Cruz Biotechnology, CTCF (07-729) from Millipore, Rad21 (ab992), H3 (ab1791) and H3K27ac (ab4729) from Abcam. Normal rabbit IgG (12-370) from Millipore was used as a negative control.

### Chromosome conformation capture (3C)

Nuclei were prepared from approximately 2 × 10^6^ of formaldehyde cross-linked cells and were digested with 600 units of Hind III restriction enzyme overnight at 37°C with shaking at 900 rpm. Fragmented DNA was ligated using 400 units of T4 ligase for 4 h at 16°C and purified by phenol extraction and ethanol precipitation. The 3C products were amplified by qPCR using SYBR green. Control templates were prepared as described previously (Kang *et al.,* 2017) and compared with 3C products to correct ligation efficiency between fragments and PCR efficiency of primer sets. Sequences of primers for 3C assay are presented in Supplementary Table S4.

### Trichostatin A treatment and western blot

MEL/ch11 cells (5 × 10^5^ cells/ml) were treated with 25 ng/ml Trichostatin A (TSA, Sigma) for 6 h and 24 h in complete DMEM medium. Following the treatment, protein was extracted using RIPA buffer with sonication. Equal amount of protein was separated electrophoretically in 4-15% SDS-PAGE gradient gel and the gel was transferred to 0.2 μm PVDF membrane. After blocking with 5% skim milk, the membrane was incubated with the primary antibodies against H3K27ac (ab4729, 1:10,000) and H3 (ab1791, 1:10,000), and then with the secondary antibody rabbit IgG-HRP (sc-2030, 1:5,000) in PBST (0.1% tween 20) with 1% skim milk. Protein signal was visualized using the ECL detection reagent. TSA-treated cells were used in ChIP and 3C assay.

### ChIP DNA library preparation and sequencing

ChIP DNA and input DNA (10 ng for H3K27ac, 0.5 ng for CTCF and 10 ng for input) were processed using NEBNext ChIP-seq library (New England Biolabs) with manufacturer’s instructions. The ends of ChIP DNA were repaired by NEBNext Ultra II End Prep enzyme mix and ligated with NEBNext adaptors. The concentration of adaptors was 1.5 μM for H3K27ac ChIP DNA and 0.6 μM for CTCF ChIP DNA. Adaptor-ligated DNA of 200 bp size was selected using NEBNext sample purification bead and amplified using the adaptor primers (7 cycles for H3K27ac, 12 cycles for CTCF and 7 cycles for input). The fragments are purified using NEBNext sample purification beads. Final libraries were quantified with Qubit dsDNA HS assay (Invitrogen) and single-end 100 bp reads were sequenced on an Illumina NovaSeq system.

### ChIP-seq data analysis

ChIP-seq raw reads were trimmed to 36 bp using trim sequences tool. Trimmed reads were filtered to remove input read ends with poor quality values (quality score 20) and then aligned to the hg19 canonical genome using Bowtie2 (Langmead & Salzberg, 2012). Aligned BAM files were filtered by Minimum MAPQ quality score 20 and sorted by chromosomal coordinates. Potential PCR duplicates were removed from BAM files. Peak regions were identified using MACS2 providing input chromatin data as control (Feng *et al*, 2012). The peak files were used to generate heatmap and peak profile using ComputeMatrix, plotHeatmap and plotProfile tools (Ramirez *et al*, 2016). The ChIP-seq signals were visualized using Integrated Genome Browser (IGB) (Freese *et al*, 2016). All public ChIP-seq data was downloaded from Gene Expression Omnibus (GEO) database and also mapped from raw reads to ensure consistent data across samples. Public ChIP-seq data used in the study is CTCF (GSM3671075) in HUDEP2 cells, H3K27ac_1 (GSM733656), H3K27ac_2 (GSM646434), CTCF_1 (GSM749690), CTCF_2 (GSM822311), GATA-1 (GSM1003608), TAL1 (GSM935496) and p300 (GSM935401) in K562 cells and H3K27ac (GSM1383853) and CTCF (GSM749753) in HMEC cells. Control input of each ChIP-seq data was analyzed in same way. Enrichment analysis of GO_biological process and TRRUST_transcription factors (Han *et al*, 2015) was performed using genes obtained from K562 CTCF ChIA-PET data (GSM970216) analysis through Enrichr (https://amp.pharm.mssm.edu/Enrichr/) tool.

## Data availability

ChIP-seq data are available under the GEO accession numbers GSE146827 and GSE147037.

## Acknowledgements

This research was supported by Basic Science Research Program through the National Research Foundation of Korea (NRF) funded by the Ministry of Science, ICT and Future Planning (NRF-2017R1A2B4008947) and by the Ministry of Education (NRF-2017R1D1A1B03030807).

